# CMT disease 2A and demyelination decouple ATP and ROS production by axonal mitochondria

**DOI:** 10.1101/462523

**Authors:** Gerben van Hameren, Graham Campbell, Marie Deck, Jade Berthelot, Roman Chrast, Nicolas Tricaud

**Affiliations:** Institut des Neurosciences de Montpellier, INSERM U1051, Université de Montpellier, Montpellier, France; Departments of Neuroscience and Clinical Neuroscience, Karolinska Institutet, Stockholm, Sweden

**Keywords:** axonal activity, demyelination, MFN2, mitochondria, ROS

## Abstract

Mitochondria are critical for the function and maintenance of myelinated axons notably through ATP production. A by-product of this activity is reactive oxygen species (ROS), which are highly deleterious for neurons. While ROS and metabolism are involved in several neurodegenerative diseases, it is still unclear how axonal activity or myelin modulates ATP and ROS production in axonal mitochondria. We imaged and quantified mitochondrial ATP and hydrogen peroxide (H_2_O_2_) in resting or stimulated peripheral nerve myelinated axons *in vivo*, using genetically-encoded fluorescent probes, two-photon time-lapse and CARS imaging. ATP and H_2_O_2_ productions are intrinsically higher in nodes of Ranvier even in resting conditions. Axonal firing increased both ATP and H_2_O_2_ productions but with different dynamics. In neuropathic MFN2^R94Q^ mice, mimicking Charcot-Marie-Tooth 2A disease, defective mitochondria failed to upregulate ATP production following axonal activity. However, H_2_O_2_ production was dramatically sustained. Mimicking demyelinating peripheral neuropathy resulted in a reduced production of ATP while H_2_O_2_ level soared. Taken together, our results suggest that ATP and ROS productions are decoupled under neuropathic conditions, which may compromise axonal function and integrity.

## 1. Introduction

While the nervous system, and the brain in particular, represents around 2% of the body mass, it consumes up to 20% of the glucose we mobilize every day [1]. This high energy expenditure in the nervous system is firstly due to the synaptic activity that requires high amounts of adenosine tri-phosphate (ATP) [1]. A second energy demanding process is the propagation of action potentials (APs) along axons. Indeed, conduction of APs involves ion exchanges through the plasma membrane, first through voltage-gated ion channels to depolarize, then through the sodium-potassium ATPase to repolarize [2,3]. Therefore, the production of ATP in axons is crucial for repeated regeneration of APs. Both in the central nervous system (CNS) and in the peripheral nervous system (PNS), myelination significantly reduces the energy cost of AP propagation through the sequestration of the AP firing machinery at the node of Ranvier and axon initial segment [4].

Mitochondria appear to be the main source of cellular ATP and these organelles are abundant in CNS [5,6] and PNS [7] axons. Some reports indicated that mitochondria were more abundant in the node of Ranvier [8]. In addition, axonal AP activity and axo-glial junctions regulates the recruitment of mitochondria in the nodal area [9]. Therefore, a well-accepted idea is that axonal AP activity could locally stimulate ATP production in axonal mitochondria [10]. However, this nodal enrichment in mitochondria remains controversial [11] and the dynamic of local mitochondrial ATP production following AP is actually unknown.

As an intrinsic by-product of ATP production, mitochondria also produce reactive oxygen species (ROS) [12,13]. The mitochondrial electron transport chain (ETC) and NAPDH oxidases are the major sources of superoxide (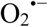), a highly reactive type of ROS, within cells [14]. 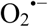 is rapidly converted to H_2_O_2_ by superoxide dismutase (SOD) and H_2_O_2_ is then reduced to H_2_O and O_2_ by mitochondrial glutathione peroxidase (GPx) and cytosolic enzymes [15]. H_2_O_2_ can also react with iron or 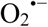 to form hydroxyl radicals (OH•), which is then reduced by molecular hydrogen to H_2_O [16]. If not reduced by antioxidants, all types of ROS are highly toxic for the cell and in particular for mitochondria, where they are produced. Indeed ROS can directly oxidize and damage DNA, carbohydrates, proteins or lipids [17].

In several neurodegenerative diseases and axonopathies, such as Parkinson’s disease [18], multiple sclerosis (MS) [19] and Charcot-Marie-Tooth diseases (CMT) [20], mitochondrial dysfunction and increased levels of ROS have been shown to be involved. Whether axonal mitochondria are involved is still unclear and investigating the production of ATP and ROS by these mitochondria in active or diseased axons in real time *in vivo* would constitute a first answer.

The importance of mitochondrial homeostasis for the normal function of myelinated axons [9,21] is highlighted by mutations in the *mitofusin 2* (*MFN2*) gene. Indeed, MFN2 has several functions in mitochondria physiology, among which mitochondrial fusion [22] and transport along axons [23], and mutations in its gene result in axonal form of peripheral neuropathy (CMT2A) in humans and in mice [24,25]. In particular, MFN2 loss-of-function in axonal mitochondria decreases the expression of oxidative phosphorylation subunits [26], suggesting that *MFN2* mutations also affect ATP and ROS production. However, this remains to be shown *in vivo*.

MS is a chronic inflammatory disease affecting myelinated tracts in the brain [27]. MS is characterized by demyelinated lesions resulting from the loss of the myelin sheath and the progressive form of MS shows successive events of myelin degeneration and regeneration that damage axons [27]. A common hypothesis is that demyelinated axons are more demanding in ATP in order to maintain their functions and their integrity [28]. Recent data showed that axonal mitochondria complex IV activity is increased in MS lesions and the total mitochondrial mass is increased [29], suggesting that axonal mitochondria are indeed producing more ATP. In addition, while activated microglia and macrophages are thought to be the initial source of ROS, axonal mitochondria dysfunction and an increase of mitochondrial ROS contribute to progressive MS [30]. Taken together this suggests that the myelin sheath plays a significant role in the maintenance of axonal mitochondria homeostasis and in particular in the production of ATP and ROS in mitochondria.

To answer these questions, we set up an *in vivo* approach using a H_2_O_2_-sensitive GFP (roGFP-Orp1) [31] and a fluorescent ATP sensor (ATeam) [32] targeted to mitochondria to image and measure the dynamics of ATP and H_2_O_2_ production in mitochondria of myelinated axons in myelinated axons of the PNS. Using this approach, we show that both ATP and H_2_O_2_ levels are increased in mitochondria residing in nodes of Ranvier and that nerve stimulation sharply increases ATP and H_2_O_2_ production by mitochondria in few minutes. While MFN2 mutations had no effect on basal levels of ATP and H_2_O_2_ in axons, it prevented the increase of ATP production after nerve stimulation. At the opposite, an increased H_2_O_2_ production was observed. Moreover, pathological demyelination reduced ATP production while enhancing mitochondrial ROS, showing that both neuropathic conditions decoupled the production of ATP from the production of ROS.

## 2. Results

### 2.1. Validation of the mito-roGFP-Orp1 and mito-ATeam probes

We used ATeam, which is a genetically-encoded fluorescence probe based on fluorescence resonance energy transfer (FRET), to detects ATP [32]. This probe consists of the cyan fluorescent protein (CFP) mseCFP, the yellow fluorescent protein (YFP) variant monomeric Venus, both linked at the ε-subunit of the Bacillus subtilis FoF1-ATP synthase. In the presence of ATP, the ε-subunit retracts to bring the two fluorescent proteins close to each other, thereby increasing FRET efficiency. The conformational change is reversible. Relative ATP levels are measured using the Venus/CFP fluorescence ratio.

Redox sensitive GFPs (roGFPs) harbor an engineered dithiol/disulfide switch on their surface, which determines their wavelength of excitation [33]. RoGFP2 has been converted into an H_2_O_2_ specific probes by fusion with the microbial H_2_O_2_ sensor Oxidant Receptor Peroxidase 1 (Orp1) [31]. The reaction is reversible via reduction by cellular thioredoxin (Trx) or GRx/GSH [34]. Relative H_2_O_2_ levels are measured using the ratio of emitted fluorescence lights when excited at 800nm (oxidized form) or 940nm (reduced form). All experiments were done in the same imaging conditions and analysis. As both probes are ratiometric, the measured values are independent of the number of mitochondria or of the absolute fluorescence intensity of the probe. Both probes were targeted to the mitochondrial matrix.

We first validated mito-roGFP-Orp1 and mito-ATeam probes *in vivo.* Injection of AAV9-mito-roGFP-Orp1 and AAV9-mito-ATeam in the spinal cord of mouse pups one day after birth (P1) resulted in expression of the fluorescent probes in multiple axons that are part of sciatic and saphenous nerves (Fig 1A). Coherent Anti-stokes Raman Scattering (CARS) microscopy uses the non-linear interactions between light and molecules to generate light from lipids without labeling [35]. As myelin is enriched with lipids, CARS microscopy allows visualizing the myelin sheath around axons *in vivo* [36,37,38]. Using this technique, we observed that the fluorescent probe signal was encompassed by the myelin sheath, which shows the probes are expressed in myelinated axons (Fig 1A). We then performed an immunostaining of teased fibers for the mitochondrial marker TOM20 and observed a partial colocalisation between fluorescent probes and mitochondria (Fig 1B), showing probes are expressed in axonal mitochondria, but not all of them. Consistently, mitochondria located in Schwann cells surrounding axons were labeled with TOM20 antibody but not with fluorescent probes. Both probes could be detected in axons of the mouse saphenous and sciatic nerves *in vivo.*

**Figure 1:**
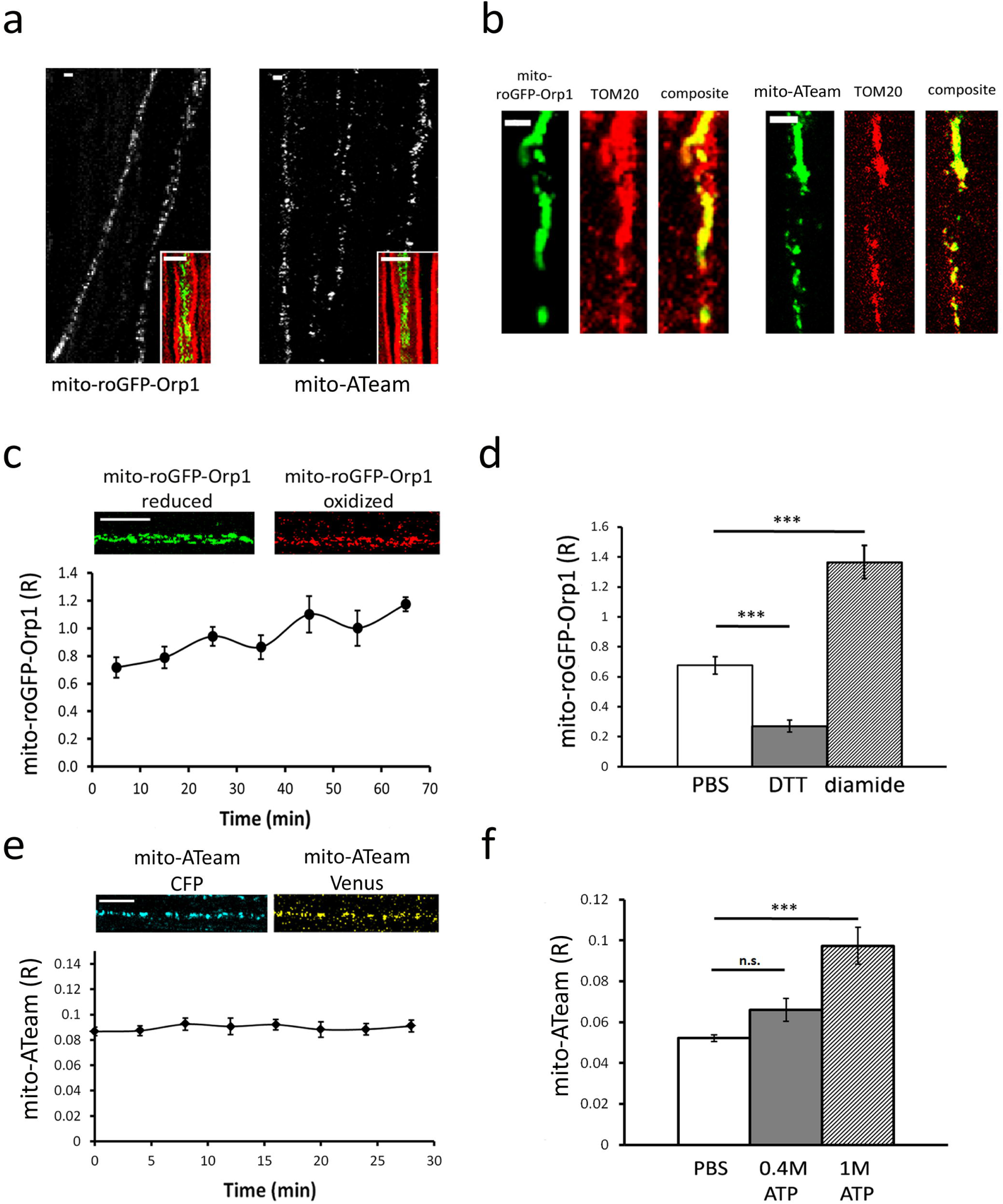
*In vivo* validation of the fluorescent probes. (A) Several axons expressing mito-roGFP-Orp1 or mito-ATeam probe can be observed in teased fibers of mouse sciatic nerve 1 month after the virus injection. CARS imaging (inserts) shows that axons expressing mito-roGFP-Orp1 or mito-ATeam (green) are surrounded by a myelin sheath (red). Scale bars = 3 μm. (B) mito-roGFP-Orp1 expression (green) partially colocalizes with mitochondrial marker TOM20 (red) in the axon. Similarly, mito-ATeam partially colocalizes with TOM20 in axonal mitochondria. Scale bars = 3 μm. (C) A fluorescent signal of mito-roGFP-Orp1 is detected when the probe is reduced and when it is oxidized by H_2_O_2_. The ratio of mito-roGFP-Orp1 fluorescence measured *in vivo* (n=6 axons; 6 mice) shows a slight increase over time. Scale bar = 10 μm (D) Mito-roGFP-Orp1 fluorescence ratio decreases with DTT (p=0.006; n=3 axons, 3 mice) and increases after injection of diamide into the nerve (p=0.003; n=4 axons, 4 mice). (E) A fluorescent signal of the CFP subunit and Venus subunit of mito-ATeam is detected. Mito-ATeam fluorescence ratio measured *in vivo* shows no significant change over time (n=18 axons; 3 mice). Scale bar = 10 μm. (F) Mito-ATeam fluorescence ratio increases after injection of 0.4M (p=0.06; n=3 axons, 3 mice) and 1M (p=7.5E-4; n=6 axons, 4 mice) ATP into the sciatic nerve. All error bars show SEM. Statistical analysis shows Student two-tailed T-tests.

Then the ratio of the fluorescent intensity of oxidized/reduced roGFP-Orp1 (R) was followed over time (Fig 1C), showing a small and slow increase of the ratio (0.206 ± 0.002 per hour) probably due to air oxidation. Addition of reducer DTT on the nerve resulted in a significant decrease of the fluorescence ratio, while injection of oxidizer diamide resulted in a significant increase of fluorescence ratio (Fig 1D). These results show that the dithiol/disulfide switch of mito-roGFP-Orp1 is an efficient and reliable indicator of relative redox changes in axonal mitochondria of mouse peripheral nerves *in vivo*. The fluorescence ratio Venus/CFP of mito-ATeam (R) was very stable over time (Fig 1E). Injection of 0.4M and 1M ATP into the sciatic nerve in an increase of ATeam fluorescence ratio (Fig 1F), showing this probe is also functional *in vivo*.

### 2.2 The effect of nerve stimulation on ATP and H_2_O_2_ production

While ATP is supposed to be critical for the firing of axons, ATP production in mitochondria of functionally active axons has never been observed *in vivo.* We used the real-time imaging approach previously described in combination with a setup for electrical nerve stimulation based on previous work [39] (Fig 2A-D). We used the saphenous nerve, which is mostly composed of sensory axons, hence its stimulation leads to only limited unwanted contraction of the paw’s muscles [40]. After recording the probe fluorescence for at least 20 minutes without stimulation, APs were induced by an electrical burst stimulation protocol including 3 bursts of 50Hz for 30 seconds spaced by a 60 second recovery period (Fig 2E).

**Figure 2:**
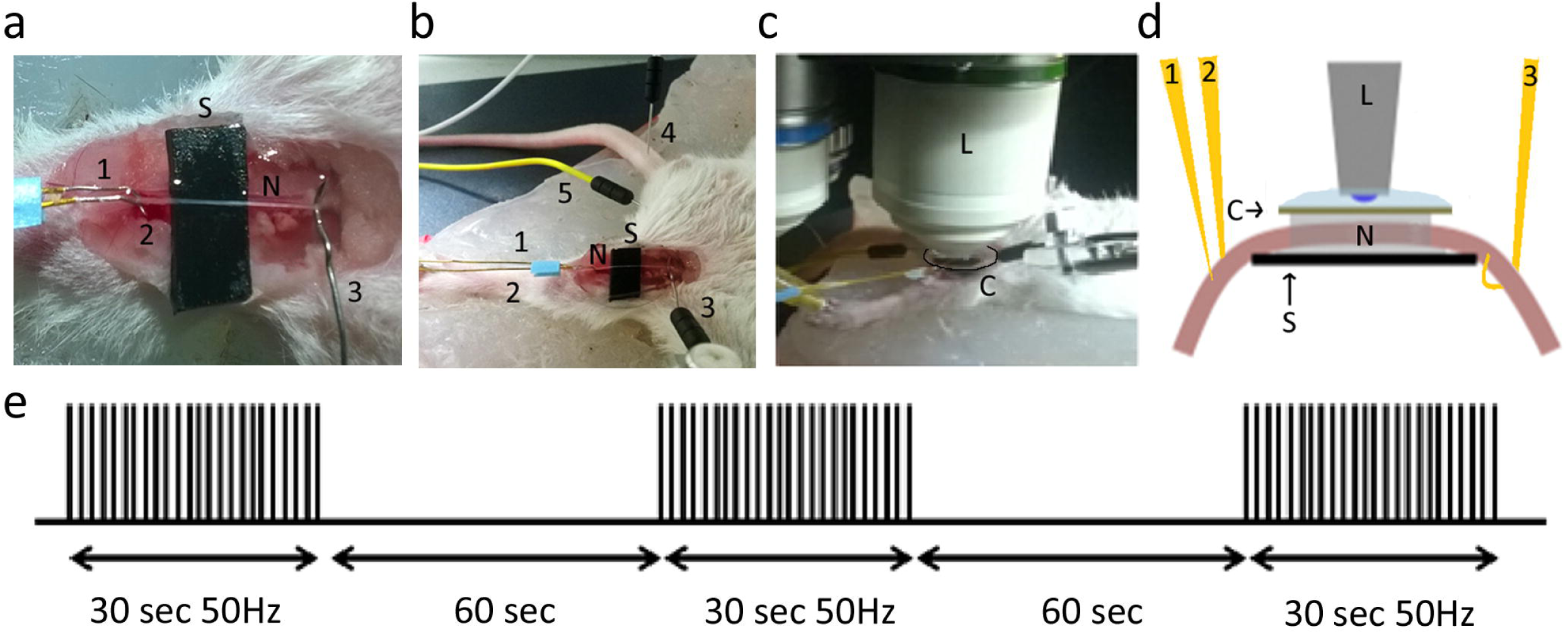
Setup of the saphenous nerve imaging experiment. (A) After deep anesthesia, the mouse is placed on its back, the skin of its left inner tight is removed and the saphenous nerve (N) is placed on a plastic strip (S). Two stimulation microelectrodes (1 and 2) are inserted on both sides of the nerve on the plastic strip and one hook-shaped recording electrode (3) is holding the nerve on the opposite side. (B) The ground electrode (4) is inserted in the mouse tail and the negative electrode (5) in the groin area. (C) A glass coverslip (Co) is then placed on the nerve soaked in PBS and the mouse is then placed under the two-photon microscope immersion lens 20X (L) in water. (D) Schematic view of the imaging setup. (E) Schematic representation of the nerve stimulation pattern used to induce APs. The generation of APs was verified using the recording electrode. Negative electrode was used to correct for background signal.

The fluorescent signal of mito-ATeam was recorded every 5 minutes for 20 minutes after stimulation (Fig 3A). At each time point the fluorescence ratio (R) was corrected for the mean fluorescence ratio of that same axon before nerve stimulation (R0). After a first stimulation, the fluorescence ratio of mito-ATeam increased in several axons, but not in all of them (44%), resulting in a slight non-significant average increase (Fig 3a). After the second stimulation, all axons responded significantly increasing mitochondrial ATP production (Fig 3a). In the same conditions, nerve stimulations significantly increased mitochondrial H_2_O_2_ production just 1 minute after the stimulation period (Fig 3B). H_2_O_2_ level then quickly went back and stabilized at pre-stimulation values (Fig 3B). These data show that axonal mitochondria very quickly adapt to the axonal activity up-regulating their production of ATP. Spikes in mitochondrial H_2_O_2_ production were also observed just before the surge of ATP in mitochondria (Fig 3C), suggesting that this H_2_O_2_ production reflects the oxidative phosphorylation process.

**Figure 3:**
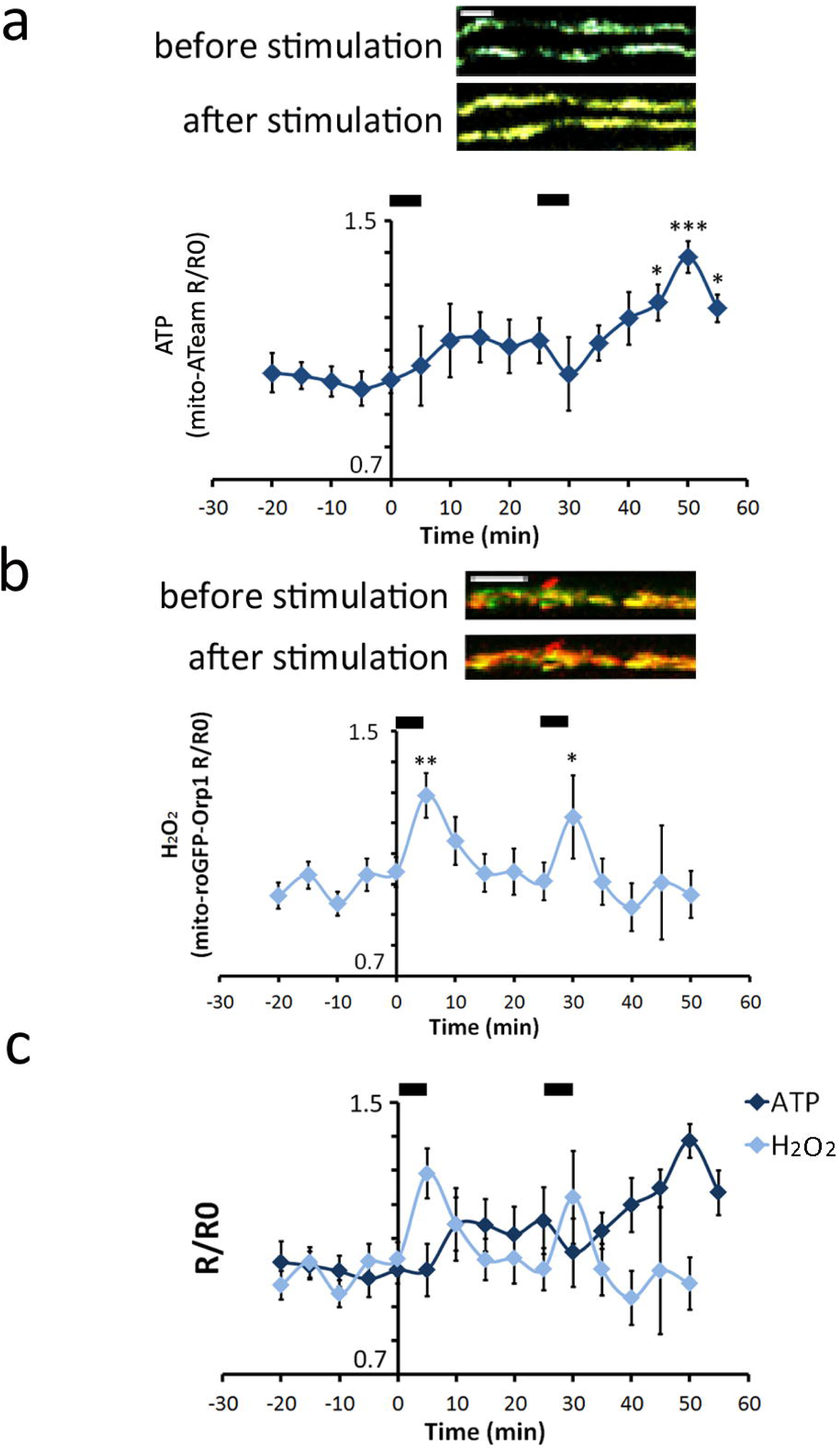
Effect of burst nerve stimulation on mitochondrial ATP and H_2_O_2_ levels. (A) Upper panels: Nerve stimulation induced changes in the fluorescence signal of both the CFP and Venus subunit of mito-ATeam as illustrated by the Venus/CFP overlay pictures. Lower panel: graph showing mito-ATeam fluorescence ratio normalized to pre-stimulation values (R/R0) following two successive nerve stimulation period (black bars at the top). The first stimulation results in a slight, non-significant increase; a significant increase is measured after a second stimulation (p=0.0171 at t=45, p=0.0001 at t=50, p=0.049 at t=55; F-value = 2.872; Df=15; n=9 axons in 3 mice). (B) Upper panels: Nerve stimulation induced changes in the fluorescence signal of both the oxidized and reduced forms of GFP in mito-roGFP-Orp1 as illustrated by the overlay pictures. Lower panel: graph showing mito-roGFP-Orp1 fluorescence ratio normalized on pre-stimulation values (R/R0) following two successive nerve stimulation (black bars at the top). Both stimulations result in a significant increase 5 minutes after the stimulation (p=0.007; p=0.04; F-value = 2.804; Df=14; n=14 axons in 7 mice). (C) Graphs described in a and b were overlaid to show the relative dynamics of ATP and ROS levels after nerve stimulations. Scale bars= 5 μm. Error bars show SEM. Statistical tests are one-way ANOVA.

### 2.3. ATP and H_2_O_2_ production are altered in CMT2A neuropathic mice

We recently showed that MFN2^R94Q^ mice, a model for CMT2A disease [25] where MFN2 is defective, displayed altered mitochondria motility and clustering in peripheral nerve axons [41]. We used this model to measure the level of ATP in axonal mitochondria of myelinated axons in neuropathic conditions. In non-stimulated conditions, control and mutant mice axonal mitochondrial ATP production were similar (Fig 4A). After the first stimulation, similar to control mice, mitochondrial ATP production increased in some axons whereas others did not respond, resulting in statistically non-significant variation (Fig 4B). However, after a second period of nerve stimulation, as opposed to controls, MFN2^R94Q^ axonal mitochondria failed to upregulate ATP production (Fig 4B-C). This indicates that dysfunctional MFN2 does not hinder the basal production of ATP by axonal mitochondria but impairs their ability to up-regulate their production in response to axonal activity.

**Figure 4:**
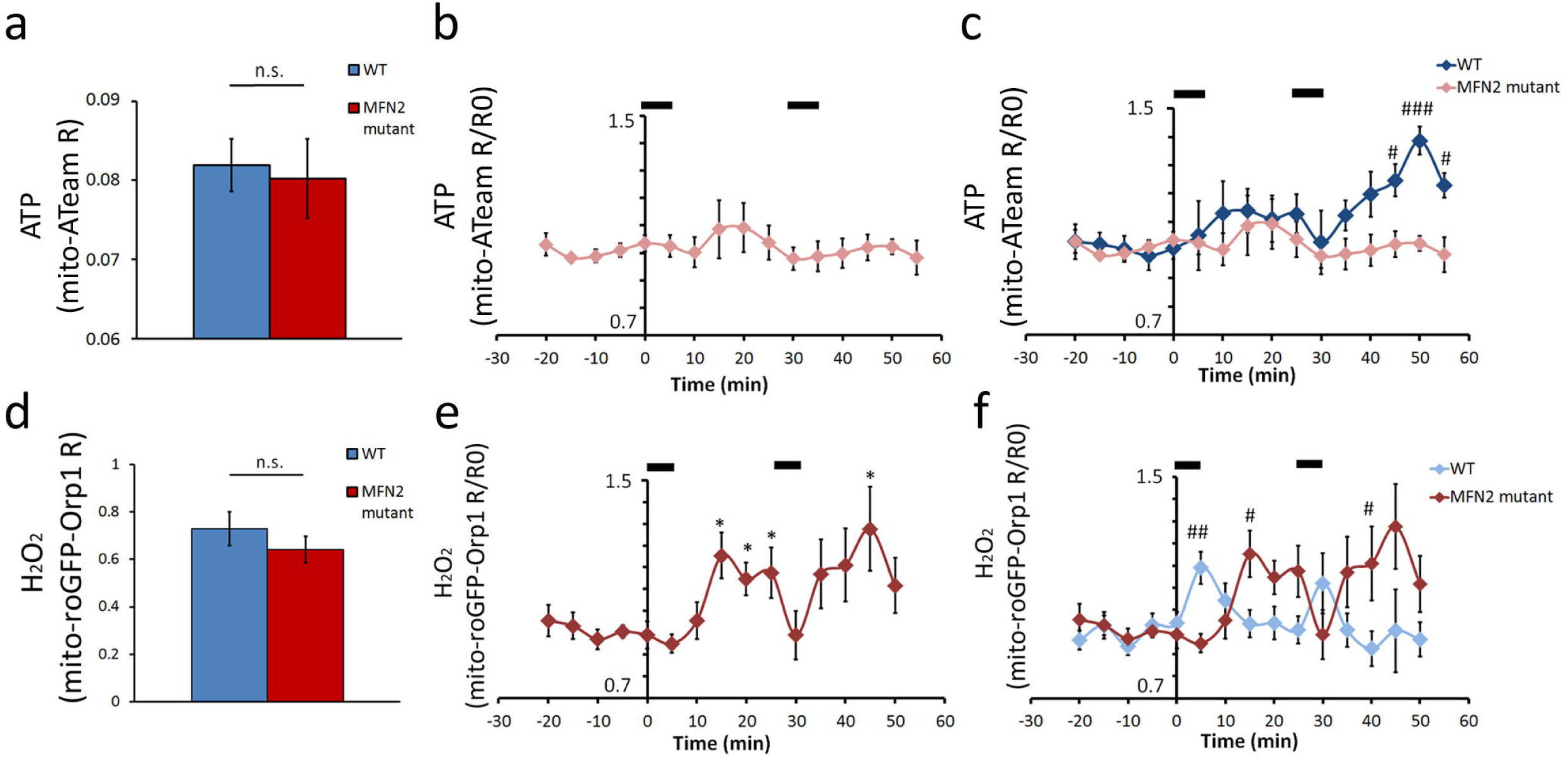
ATP and H_2_O_2_ levels of MFN2^R94Q^ mice. (A) Graph showing mito-ATeam fluorescence ratio in resting axonal mitochondria of wild-type or mutant MFN2^R94Q^ mice (Student two-tailed T-test p=0.809; n=18 axons, 7 mice per group). (B) Graph showing mito-ATeam fluorescence ratio normalized on pre-stimulation values (R/R0) following two successive nerve stimulation periods (black bars at the top). The first period of stimulation resulted in a slight, non-significant increase; the second stimulation period did not generate any change (n=14 axons in 5 mice). (C) Graphs showing mito-ATeam R/R0 in wild-type (Fig. 3A) and mutant MFN2^R94Q^ mice (B panel) were overlaid to show the relative dynamics of ATP levels in both genotypes after nerve stimulations (Two way ANOVA, p=0.011 at t=45, p=4.8E-5 at t=50, p=0.02 at t=55, F-value=19,27; Df=1). (D) Graph showing mito-roGFP-Orp1 fluorescence ratio in resting axonal mitochondria of wild-type or mutant MFN2^R94Q^ mice (Student two-tailed T-test p=0.337; n=16 axons, 8 mice per group). (E) Graph showing mito-roGFP-Orp1 fluorescence ratio normalized on pre-stimulation values (R/R0) following two successive nerve stimulation periods (black bars at the top). Both stimulations result in a significant increase (p=0.013 at t=15, p=0.020 at t=20, p=0.044 at t=25 and p=0.038 at t=45; F-value=3.095; Df=14; n=11 axons in 4 mice). (F) Graphs showing mito-roGFP-Orp1 R/R0 in wild-type (Fig. 3B) and mutant MFN2^R94Q^ mice (E panel) were overlaid to show the relative dynamics of H_2_O_2_ levels in both genotypes after nerve stimulations. A significant higher H_2_O_2_ level is detected in wild-type first (Two way ANOVA, p=0.007 at t=5, F-value= 5,484; Df=1), then a higher H_2_O_2_ level is detected in mutant MFN2^R94Q^ (p=0.014 at t=15, p=0.047 at t=40). Error bars show SEM. # = p-value<0.05. ## = p-value<0.01. ### = p-value<0.001.

Similar to ATP, we found no difference between controls and MFN2^R94Q^ mice in the basal H_2_O_2_ levels (Fig 4D). When nerves were stimulated, mitochondrial H_2_O_2_ significantly increased after each stimulation period (Fig 4E). However, this increase was delayed compared to controls and H_2_O_2_ levels remained high for longer time (Fig 4F), showing that axonal mitochondria produce more deleterious H_2_O_2_ than control mice in response to axonal activity. Since ROS production in the mitochondrial matrix reflects the oxidative phosphorylation process [42,43], these data indicate that oxidative phosphorylation is decoupled from ATP production in mitochondria of these neuropathic mice.

### 2.4. ATP and H2O2 production in nodes of Ranvier and internodes

The enrichment of mitochondria at the node of Ranvier and whether these nodal mitochondria are more metabolically active remains controversial. In order to detect potential spatial differences between nodal and internodal mitochondria, we used CARS microscopy. Gaps in CARS signal between two myelinated internodes are nodes of Ranvier (Fig 5A) [38,43,44]. Combining two-photon imaging of fluorescent probes with CARS imaging allowed us to analyze mitochondrial physiology in nodes of Ranvier versus internodes defined as areas distant of more than 5 μm from the node (Fig 5A). In all observed nodes of Ranvier, probes-labeled mitochondria were present, but they were not more abundant or larger than in internodes (Fig 5A). However, when measuring the mito-ATeam and mito-roGFP-Orp1 fluorescence ratio, we found significantly higher ATP and H_2_O_2_ levels in node of Ranvier mitochondria (Fig 5B-C). These data, obtained in resting animal without nerve stimulations, indicate that nodal mitochondria are intrinsically more metabolically active than internodal ones. In neuropathic MFN2^R94Q^ mice, nodal mitochondria did not show a higher level of ATP versus internodal mitochondria (Fig 5B). However, they still had higher level of H_2_O_2_ (Fig 5C), suggesting again a decoupling between oxidative phosphorylation and the production of ATP in mitochondria of these mutant mice.

**Figure 5:**
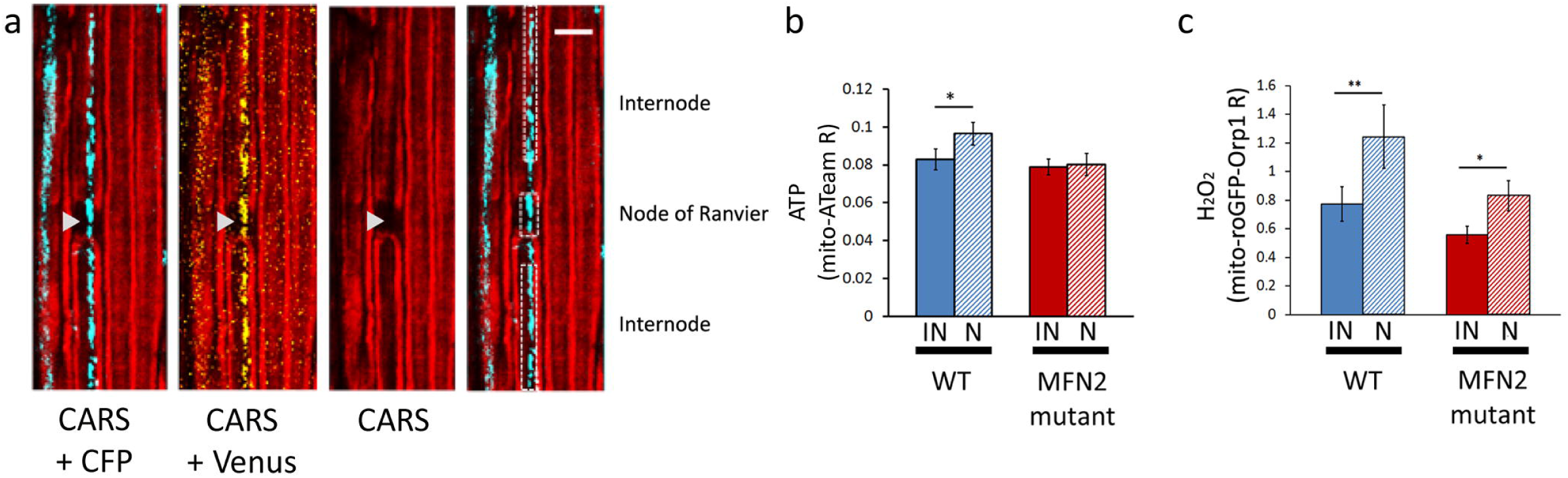
ATP and H_2_O_2_ levels are different along axons. (A) Two-photons imaging of CFP (blue) and Venus (yellow) fluorescence of mito-ATeam was combined with CARS imaging of the myelin sheath (red). CARS imaging gap shows a node of Ranvier (arrowhead). Using the combined images, mitochondria located in the node of Ranvier and mitochondria located in internodes can be identified (right panel). Scale bar = 10 μm (B) Graph showing mito-ATeam fluorescence ratio in axonal mitochondria located in nodes of Ranvier (N) or internodes (IN) of wild-type (WT) or MFN2^R94Q^ mice (MFN2 mutant). In wild-type mice, ATP levels are higher in mitochondria in nodes of Ranvier than in internodes (p=0.035; n=14 axons in 7 mice). In MFN2^R94Q^ mice, ATP levels in node of Ranvier mitochondria are equal to internode mitochondria (n=7 nodes in 3 mice). (C) Graph showing mito-roGFP-Orp1 fluorescence ratio in axonal mitochondria located in nodes of Ranvier or internodes of wild-type or mutant MFN2^R94Q^ mice. In both wild-type mice and mutant MFN2^R94Q^ mice, H_2_O_2_ levels are higher in mitochondria in nodes of Ranvier than in internodes (WT: p=0.009; n=8 axons in 3 mice. MFN2^R94Q^: p=0.019; n=9 axons in 3 mice). Error bars show SEM. Statistical tests are paired two-sided T-tests.

### 2.5. ATP and H_2_O_2_ production in mitochondria of demyelinated axons

The myelin sheath is critical to maintain the node of Ranvier on myelinated axons and is therefore likely to play a significant role in the homeostasis of axonal mitochondria. As demonstrated in Fig 5a, the myelination state of the axons in the mouse sciatic nerve can be assessed *in vivo* by CARS imaging. We therefore investigated how this demyelination affects axonal mitochondria physiology *in vivo.* To induce demyelination we used lyso-phosphatidylcholine (LPC) and we followed the production of mitochondrial ATP and H_2_O_2_ in axons during demyelination and during the restoration of the myelin sheath (remyelination). LPC, which is a signaling molecule resulting from the hydrolysis of phophatidylcholine by phospholipase A2 [45] acts in mSC and oligodendrocytes as a demyelinating signal [46,47,48]. LPC was injected in the sciatic nerve of adult mice and demyelination occurred immediately to culminate one week after the injection with the formation of myelin ovoids and debris as seen using CARS (Fig 6A, week 1) [37,49]. As soon as two weeks after LPC injection, remyelination occurred and some thin myelin sheaths reappeared around axons (Fig 6A, week 2). Axons were completely remyelinated three weeks after injection of LPC (Fig 6a, week 3). No demyelination occurred in control mice injected with PBS (Fig 6A last panel). Axonal integrity is not directly affected by LPC (S1 Figure) [50] and axonal mitochondria remained visible using either mito-ATeam or mito-roGFP-Orp1 probes during the entire process of demyelination and remyelination (Fig 6A inserts).

**Figure 6:**
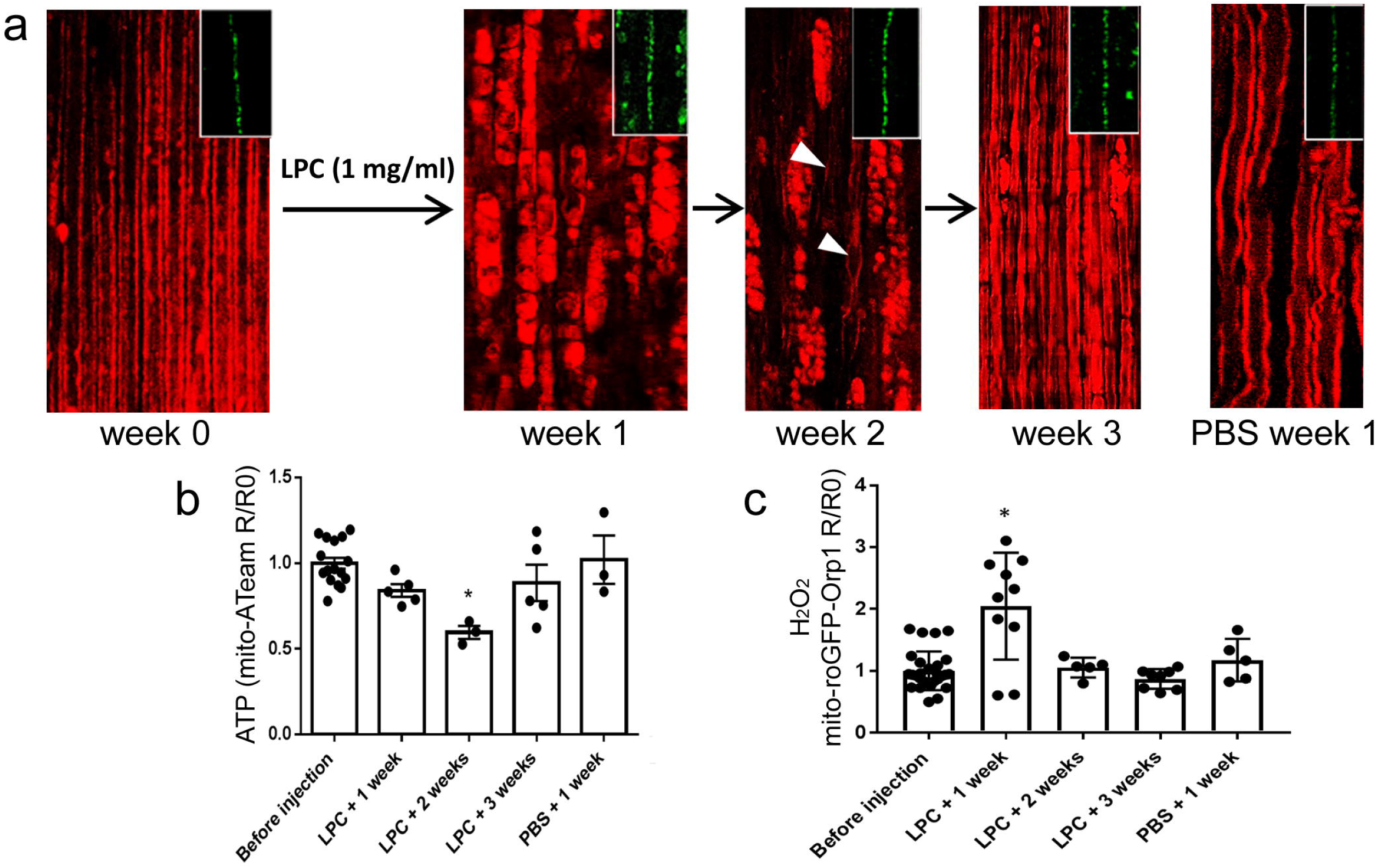
Impact of demyelination on axonal mitochondria ATP and H_2_O_2_. (A) CARS imaging is used to visualize myelin. At week 0, before injection of LPC, most axons are myelinated. 1 week after LPC injection, axons are demyelinated and myelin debris and ovoids are observed. At week 2, thinly myelinated axons are observed in between the myelin debris (arrowheads) and at week 3, almost all axons are myelinated again, thus resembling a healthy nerve. PBS injection induces small conformational changes, but no formation of ovoids or debris. Inserts show that neuronal mitochondria remain visible throughout the whole process. (B) Graph showing mito-ATeam fluorescence ratio (R) in axonal mitochondria normalized on pre-demyelination values (R0) following demyelination. Following LPC injection, ATP levels are unchanged after 1 week (p=0.085; n=5 mice; 27 axons), decreased after 2 weeks (p=0.019; n=3 mice, 24 axons) and restored after 3 weeks (p=0.491; n=5 mice; 31 axons). PBS injection results in no significant change (p=0.799; n=3 mice; 23 axons). (C) Graph showing mito-roGFP-Orp1 fluorescence ratio (R) in axonal mitochondria normalized on pre-demyelination values (R0) following demyelination. Following LPC injection, H_2_O_2_ levels are increased after 1 week (p=0.031; n=10 mice, 17 axons), and unchanged after 2 weeks (p=0.663; n=5 mice, 9 axons) and after 3 weeks (p=0.450; n=8 mice, 13 axons). PBS injection results in no significant change (p=0.121; n=3 mice, 5 axons).

We therefore measured the fluorescence ratio for these probes in non-stimulated axonal mitochondria during the process. One week after LPC injection, when axons are completely demyelinated, ATP production was slightly lower than before demyelination, but not significantly (Fig 6B), while H_2_O_2_ levels peaked (Fig 6C). ATP levels were significantly lower in axonal mitochondria at two weeks after LPC injection, while H_2_O_2_ levels were back to control and pre-demyelination levels (Fig 6B-C). Finally, three weeks after injection, when remyelination is completed, ATP levels were back to control and pre-demyelination levels as are the H_2_O_2_ levels (Fig 6B-C). No change either in ATP or H_2_O_2_ levels were detected at one week after injection with PBS in control mice (Fig 6B-C). Moreover, increase in ROS did not result from a general oxidative stress in the demyelinated nerve, since CellRox dye did not show any significant increase, except at the LPC injection site where we did not image our probe (S1 Fig). Taken together, these data indicate that the myelin sheath has a profound impact on axonal mitochondria physiology. Mitochondria manage to maintain their ATP production during myelin break-down, but they generate more H_2_O_2_. The process of remyelination leads to decreased production of ROS, while ATP production is temporally diminished before full remyelination. These data therefore indicate that demyelination induces the decoupling of ATP production from oxidative phosphorylation.

## 3. Discussion

Although it has been shown that PNS axonal mitochondria, like most mitochondria, produce both ATP and ROS, the regulations of their production are still not well understood *in vivo* and *in situ*. We used a combination of viral delivery of mitochondria-targeted fluorescent probes to active peripheral neurons and two-photon and CARS live imaging in mice to fill that gap. The mito-ATeam and mito-roGFP-Orp1 fluorescent probes had been previously validated *in vitro* [51] and *in vivo* [33] in other systems but we also validated them *in vivo* in our system. We did not attempt to measure absolute values of ATP or H_2_O_2_ with each probe fluorescence *in vivo* because firstly, this was difficult as probes are targeted to mitochondrial matrix and secondly, it was not necessary as we aimed to compare different genotypes and conditions in particular with time. So only relative values obtained in the same imaging conditions are shown here. We found that the range of detection was large enough to cover the changes occurring in axonal mitochondria *in vivo*. Moreover the probe sensitivity was sufficient to observe variations in real-time using a 5 minutes delay between measures. This delay was partly imposed by relative low speed of our imaging system (1 minute to create 10 scans of 200μm^2^ over 40 μm depth for each wavelength), but also by the intrinsic limits of our set-up: electrical stimulation induced slight contractions of saphenous nerve surrounding muscles which made the pictures blurred during the stimulation. The probes alterations were also reversible fast enough to observe changes in both directions. Mito-ATeam was stable, but mito-roGFP-Orp1 showed a slight increase of its fluorescence ratio over time under our experimental settings. These results are not explained by photobleaching or degradation of the probe, since the total fluorescent signal does not decrease. This change in ratio of fluorescence may be caused by air oxidation [52]. However, buffered artificial cerebrospinal fluid as an incubation medium for the saphenous nerve did not prevent the probe oxidation. Nonetheless, this slow oxidation of mito-roGFP-Orp1 had negligible effects on our experiments, since it was too slow to cause significant changes in our relatively short term imaging.

Axonal activity induced stereotypic changes in ATP and H_2_O_2_ production by axonal mitochondria. Shortly after the stimulation, H_2_O_2_ production increased and dropped immediately and then ATP increased for several minutes. This was stereotypic, because it occurred almost identically after both stimulations we did. After the first stimulation, not all axonal mitochondria produced more ATP, suggesting that either our stimulation protocol was not mobilizing enough axons or axonal mitochondria are deactivated, similar to some ion channels. This limitation was lifted by the second stimulation. On the attempts we did for further stimulations, sometimes the ATP level was maintained at high range and sometimes mitochondria did not respond anymore. The reason for these variations in the long term is unknown and we did not pursue this further. However, taken together, this suggests that ATP production in axonal mitochondria is positively correlated with the level of axonal activity and can eventually reach a plateau in our conditions. Due to technical reasons we could not measure H_2_O_2_ levels in mitochondria after several stimulations. However, looking at the first two stimulations, we conclude that H_2_O_2_ production in mitochondria is also positively correlated with axonal activity but not cumulative. It would be interesting to look further for the molecular mechanisms that allow mitochondria to detect axolemma depolarization or ion channels activity on a very short period of time (around one minute in our detection capacities). This fast kinetic suggests the role of ions, such as calcium ions, which are mobilized during AP firing at the node of Ranvier [53], or kinases. A mechanism involving calcium has been shown to anchor mitochondria at the node of Ranvier [54], suggesting that mitochondria are associated with the axolemma there. Consistently, we found nodal mitochondria to be more metabolically active than internodal ones. However, our data following stimulations were recorded on mitochondria independently of their location on axons. As the node is much smaller than the internodes (1 μm versus 600 μm in average in the mouse nerve) the probability we recorded nodal mitochondria is extremely small. So the molecular mechanism that mobilizes mitochondria after AP firing has to exist also in internodal mitochondria.

Looking at the known origin of ROS in mitochondria, our data suggest that the H_2_O_2_ production we detected in axonal mitochondria reflects the loading of the mitochondrial matrix with protons before the production of ATP. However, since the H_2_O_2_ production drops very quickly and, contrary to the ATP production, it is not cumulative, a more complex scheme has to be taken in consideration. Indeed, H_2_O_2_ is formed by SOD enzyme from 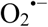, which directly results from the ETC. Moreover, H_2_O_2_ is then changed in H_2_O and O2 by several reducing enzymes, such as mitochondrial GPx, so actually the amount of H_2_O_2_ we detected is the result of equilibrium between 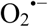 production and SOD and reducing enzymes’ activities. As these enzymes are highly expressed in mitochondria [55], they probably do not constitute a limiting factor and the H_2_O_2_ levels we detected still reflect axonal mitochondria metabolic activation. However, H_2_O_2_ levels drop quickly, probably because reducing enzymes are removing it quickly. In addition, these enzymes’ activities change following fast post-translational modifications such as phosphorylation [56] or environmental changes [57]. While this may explain why H_2_O_2_ levels are not incremental following several stimulations, further investigation will be needed to clarify underlying mechanisms.

Several CNS and PNS neuropathies are linked to mitochondrial dysfunctions in neurons [18,19,20]. Our data show that the electrical activity of neurons directly regulates the physiology and the respiration of axonal mitochondria. As axonal mitochondria are the most abundant among neuronal mitochondria, this suggests that axonal mitochondria dysfunctions may severely impact neuronal physiology and survival. We evaluated this hypothesis using a couple of peripheral neuropathy mouse model, such as a model of CMT2A disease expressing mutant MFN2^R94Q^ [25]. While *in vitro* data had shown increased H_2_O_2_ levels [58], but no change in ATP production [59] in neuronal mitochondria of these mice, *in vivo* we did not measure significant differences in these parameters between resting MFN2^R94Q^ mice and control mice. A first explanation for this discrepancy is that cultured cells are living in an environment that is significantly different from the *in vivo* conditions. This is especially true in terms of metabolism [60]. Another explanation is that mitochondria physiology is altered, but this can only be seen when mitochondria are challenged. Environmental changes in culture may constitute this challenge, while more physiologically relevant challenges have to be found *in vivo*. Our electrical stimulations were physiologically meaningful challenges for axons and, similar to control mice, we observed ATP levels increasing in several axons of mutant mice, but not all, after a first nerve stimulation. However, while control mice mitochondria increased their ATP production further following a second stimulation, no increase was observed in mutant mice mitochondria. We have shown that Mitochondria Associated Membranes (MAM), which mediate interactions between the endoplasmic reticulum and mitochondria [41], are impaired in these mutant mice. So we suggest that an impaired Ca^2+^ uptake by mitochondria through MAM [61] could alter the ATP production as it has been shown that calcium is required for the TCA cycle [62]. However, mutant mice mitochondria were still able to produce a large amount of H_2_O_2_ following both stimulations, suggesting the ETC and therefore the TCA cycle are working properly. Yet this H_2_O_2_ production was completely abnormal: it was largely delayed and was sustained for over at least 10 minutes. As discussed before the levels of H_2_O_2_ result from an equilibrium between several parameters, so it is difficult to explain this abnormal production. Nevertheless, a relationship between mitochondrial shape and ROS has been shown *in vitro* [63]. It has been suggested that smaller mitochondria result in intra-mitochondrial redistribution of cytochrome c and the pro-aging molecule p66shc that acts trough ROS upregulation [64,65]. Smaller fragmented mitochondria, such as MFN2 mutants [63], correlate with a malfunctioning antioxidant system and an increased production of ROS [58,64]. In any case, our data show that an altered MFN2 function lead to the dysfunction of axonal mitochondria only when axons are firing. This dysfunction is strongly deleterious as mitochondria don’t produce more ATP but show a sustained production of deleterious ROS. This functional decoupling between ATP and H_2_O_2_ production, occurring in the most abundant mitochondria of the neurons and in conditions of normal firing activity, is likely to be a significant cause of the neuronal dysfunction in this peripheral neuropathy.

Looking at the node of Ranvier of MFN2 mutant mice, we found that, similar to control mice, nodal mitochondria were more metabolically active than internodal ones. As MFN2 mutant mitochondria do not increase ATP production in active axons, this suggests that nodal mitochondria are intrinsically more active even in absence of firing. This is consistent with the fact that motile mitochondria slow down in the nodes of Ranvier [6] and that they have a different morphology [66]. Moreover, nodal mitochondria may participate to axonal degeneration since this process starts in nodes of Ranvier due to high ROS levels [67]. It would have been interesting to investigate changes occurring in ATP and H_2_O_2_ production of nodal mitochondria after electrical stimulation but this was impossible due to the small size of the node and the change of focus that occurs following stimulations.

The myelin sheath that covers axons appears to be an important regulator of axonal metabolism and of mitochondria physiology. Indeed in MS, demyelination alters axonal mitochondria [19,29,68] and we show here that nodes of Ranvier devoid of myelin house more metabolically active mitochondria. To go further, we therefore investigated the impact of the loss of myelin on axonal mitochondria. LPC was the right tool because it induced demyelination without destroying axons [46,47,48]. LPC-induced demyelination resulted in a decrease of ATP levels in axonal mitochondria. This decrease was strongest after two weeks and was reversed by 3 weeks, showing that ATP production within neurons is impaired until complete remyelination. This decrease was unexpected, since AP propagation in demyelinated axons is more energy demanding: the nodal machinery and the Na^+^/K^+^ ATPase are diffused all along the axolemma and all the energy-effective benefits of myelination are lost [69]. In demyelinated area of brains affected by MS, an upregulation of ETC complexes protein expression and more complexes I and IV activity [68] was observed. Moreover, demyelinated axons showed more mitochondria [29], which is consistent with a higher energy demand.

Nevertheless, H_2_O_2_ levels strongly increased in axonal mitochondria of fully demyelinated axons in absence of general oxidative stress in the tissue. This decoupling of mitochondrial oxidative phosphorylation and ATP production is in accordance with previous studies that reported a decreased ATP production, together with increased ROS levels in demyelinating diseases [70]. This suggests that actually, mitochondrial ETC complexes are more active and produce more ROS, but this does not lead to more ATP production. Such a mechanism has been shown in particular conditions where the goal of mitochondrial activity is to produce heat. Uncoupling proteins such as UCP2 permeabilize the inner mitochondrial membrane to dissipate the proton gradient [71]. This may not be the sole utility of uncoupling as this process has been shown to occur in several neurodegenerative diseases [72]. However, so far we failed to detect unusual UCP2 expression in demyelinated nerves. The high production of H_2_O_2_ during demyelination is probably not without consequences as ROS can alter the function of the ATP synthase [73]. This may explain why ATP production remains weak in mitochondria of axons that are being remyelinated two weeks after LPC injection, despite that ROS production is back to normal. This suggests that during demyelination, axons may recruit more mitochondria [29], or use another metabolism, such as aerobic glycolysis, in order to cover their need in ATP.

While very little amount of data exists on the homeostasis of axonal mitochondria in demyelinated axons of the PNS, in the CNS, demyelination in progressive MS results in axonal mitochondria dysfunction and ROS increase [74]. The data we collected in PNS myelinated axons on ROS production by axonal mitochondria are consistent with this. This suggests that, despite the large difference that exists between PNS and CNS myelinating glia, their similar role in segregating axonal firing machinery at nodes of Ranvier is essential for mitochondria homeostasis in axons. Nevertheless, the electrical isolation of the axon and the formation of the node of Ranvier is not the only function of the myelin sheath as it also largely participates to the metabolic homeostasis of the axon through the lactate shuttle process [75,76]. This is particularly true for the CNS myelin [76]. In addition, MS disease is significantly different from peripheral nerve diseases, in particular because of the important role played by CNS glial cells astrocytes and microglia. So, it would be essential to confirm the data we obtained in PNS myelinated axons in CNS myelinated axons *in vivo*, in order to definitely conclude on the alteration of ATP and ROS production by axonal mitochondria in demyelinated lesion of progressive MS.

To conclude, combining several novel techniques such as two-photon and CARS non-linear live imaging coupled to electrical nerve stimulation and genetically-encoded fluorescent probes delivered by AAV vectors, we were able to observe the production of ATP and H_2_O_2_ by axonal mitochondria in real time and in living and active axons of mouse peripheral nerves. We demonstrated that mitochondria are more metabolically active at the node of Ranvier and we characterized the role of AP firing in the dynamics of mitochondrial ATP and H_2_O_2_ production. Moreover we show the deleterious alterations that occur in axonal mitochondria physiology of two mouse models of neuropathy, peripheral axonal CMT2A and demyelinating peripheral neuropathies. Together, our data thus provide new insight into the role of axonal mitochondria under physiological and pathological conditions.

## 4. Materials and methods

### 4.1. Cloning

pLPCX mito-roGFP-Orp1 (Addgene; #64992) was digested with XhoI/ClaI, blunted and cloned into pAAV-MCS (Cell Biolabs, Inc.) under the control of a CMV promoter. Clones were validated by sequencing. Likewise, pcDNA-mito-ATeam (from H. Imamura, Tokyo, Japan) was digested with XhoI/HindIII, blunted and cloned into the CMV promoter controlled pAAV-MCS. Mitochondria-targeting tags were CoxVIII for roGFP-Orp1 and two tandem copies of CoxVIII for mito-ATeam. AAV viral particles were produced at the viral vector production centre at the Centre de Biotecnologia Animal I Teràpia Gènica in Barcelona, Spain or the University of Nantes, France.

### 4.2. Mouse strains

Either Swiss mice (Janvier, France) or MFN2^R94Q^ mice [25] were used in the performed experiments. Mice were kept in the animal facility of the Institute for Neurosciences of Montpellier in clear plastic boxes and subjected to standard light cycles. All animal experiments were conducted in accordance with the French Institutional and National Regulation CEEA-LR-11032.

### 4.3. In vivo virus injection in spinal cord

A thin borosilicate glass capillary (Harvard Apparatus, Ref. 30-0016) was pulled with a Vertical Micropipette Puller (Sutter Instruments, P30-682) to form a glass needle. The glass needle was filled with viral solution (1 μl) and a 1 day old mouse is restrained in a position that exposes the lower back. The needle is injected through the skin using a micromanipulator and introduced into the spinal cord. The viral solution was injected over 2 minutes with short pressure pulses using a Picopump (World Precision Instrument) coupled to a pulse generator. After injection, the injection site is cleaned with betadine solution (Vetoquinol, cat. no. 3042413) for disinfection and the pup is placed back with its mother and littermates.

### 4.4. Immunohistochemistry

One month after the viral injection, the sciatic nerve was dissected and fixed in Zamboni's fixative [77] for 10 min at room temperature. After fixation, the dissected sciatic nerves are washed in PBS and incubated in successive glycerol baths (15, 45, 60, 66% in PBS) for 18 to 24 h each before freezing at –20 °C. The nerves were cut in small pieces in 66% glycerol and the epineurium sheaths removed. Small bundles of fibers were teased in double-distilled water on Superfrost slides and dried for 3 hours at room temperature. Some nerves were frozen in O.C.T. Compound (Tissue-Tek, Ref. 4583) and longitudinal sections were cut using a cryostat (Leica Biosystems, CM3050). For immunostaining, the teased fibers or longitudinal sections were incubated for 1 h at room temperature in blocking solution (10% goat serum and 0.3% TritonX100 in PBS). Then, the samples were incubated with TOM20 (FL-145) primary rabbit antibody (1/500, Santa Cruz, Ref. sc-11415), mouse anti-Myelin Basic Protein (1/500, Merck, Ref. NE1019) or rabbit NF-200 (1/500, Sigma, Ref. N4142) in blocking solution overnight at 4 °C. The next day, the samples were washed in PBS and incubated for 1h at room temperature with secondary donkey antibodies coupled to Alexa568 (1/1000, Invitrogen, Ref. A10042) or Alexa488 (1/1000, Invitrogen, Ref. A21202). Finally the samples were washed in PBS and mounted in Immu-mount (Thermo Scientific). Images were acquired at room temperature using a 20× or 40× objective, a Zeiss confocal microscope LSM710, and its associated software.

### 4.5. Imaging of sciatic nerve in living mice

One month after the viral injection, the mice were anesthetized with a constant flow (1.5 l/min) of oxygen + 5% of isoflurane in an anesthesia box (World Precision Instruments, Ref. EZ-B800) for 5 min. Thereafter the anesthesia was maintained with a mask delivering 2% isoflurane at 0.8 l/min. The eyes were protected by eye protection gel (Ocry-gel, TVM, cat. no. 48026T613/3). Intraperitoneal injection of 0.1 mg/kg buprenorphine was used for pre-surgery analgesia. The mouse was placed in a silicone mold, lying on its belly, shaved on his hind paw and the paws immobilized using small pins. The incision area was disinfected with betadine solution (Vetoquinol, cat. no. 3042413). The skin was cut using scissors, fat tissue removed, the gluteus superficialis and biceps femoris muscles were separated to reveal a cavity crossed by the sciatic nerve, and the sciatic nerve was lifted up using a small spatula. A long plastic strip was placed underneath the sciatic nerve and this strip fixed using magnets. The sciatic nerve was kept in an aqueous environment of either sterile PBS buffer or artificial cerebrospinal fluid (148 mM NaCl, 3 mM KCl, 1.4 mM CaCl_2_•2H_2_O, 0.8 mM MgCl_2_•6H_2_O, 0.2 mM NaPO_4_•H_2_O in sterile H_2_O) to prevent drying. At that point the mouse was placed under the two-photon microscope objective lens, a glass, 12mm diameter, 0.25mm thick microscope coverslip put on top of the nerve and a drop of deionized water placed on it to immerse the objective lens (20X, Carl Zeiss Microscopy, LD CApochromat, Ref. 421887-9970).

### 4.6. Demyelination procedure

After imaging of the sciatic nerve in vivo for at least 30 minutes, 5 μl of 1 mg/ml lysophotphatidyl choline (LPC) was injected directly into the sciatic nerve using a Hamilton syringe (Ref. 80930) to induce demyelination. Injection of 5 μl PBS is used as a negative control. After injection, the nerve was placed back to its original location in the body. The skin of the incision was realigned together using a blunt scalpel and stapled along the wound with two clips (Fine Science Tools; 12020-00). The area around the wound was disinfected again with betadine solution. The anesthesia mask was removed and the mouse was monitored until it had woken up. Motor function of the injected and non-injected hindlimb was followed by lifting the animal by the tail. At 1 week after LPC injection, the non-injected hindlimb showed a normal postural reflex characterized by spreading of leg whereas the injected hindlimb showed abnormal reflexes such as tremors, clasping or retracting of the paw. This behavior has been described and linked to deficient myelination in the PNS in previous studies [78]. The severity of abnormal hindlimb function decreased at 2 weeks after LPC injection and no difference in leg function between the injected and non-injected hindlimb could be observed anymore at 3 weeks after LPC injection.

### 4.7. Saphenous nerve stimulation in living mice

After skin incision and removal of connecting tissue, the saphenous nerve is lifted up and isolated using a plastic strip (Fig 2a). To stimulate the saphenous nerve, two platinum kapton microelectrodes (World Precision Instruments, PTM23B05KT) were inserted at the posterior side of the plastic strip using a micromanipulator (Fig 2A). One hook-shaped recording electrode (AD Instruments, MLA 1203) was placed at the anterior side (Fig 2A). The ground electrode was inserted in the tail of the mouse and the negative electrode in the groin area (Fig 2B). The mouse was placed under the two-photon microscope (Fig 2C), a microscope glass coverslip was placed on top of the nerve and the electrodes were connected to a Powerlab 26T (AD Instruments; ML4856). A drop of deionized water was then placed on top of the microscope glass to immerse the 20× objective lens.

### 4.8. Two-photon image acquisition

All in vivo images were obtained with a two-photon microscope LSM 7 MP OPO (Zeiss, France) coupled to a dark microscope incubator (L S1 Dark, Zeiss) in which the temperature was maintained at 37 °C (Heating Unit XL S, Zeiss, France). Mitochondria images were acquired by time-lapse recording varying from one image every minute to one image every 5 minutes during 1 hour. Each image is a stack at Maximum Intensity (ZEN software, Zeiss) of 10 scans over 40 μm depth. For ATeam imaging, a single track at 850 nm excitation wavelength is used to obtain both the CFP (em. 475 nm) and Venus (em. 527 nm) image at the same time point. For roGFP imaging, the two images were acquired for each time point using alternating tracks at 940nm and 800nm. Change of track was set after each stack. For Coherent Anti-Stokes Raman Scattering (CARS) imaging, 2 synchronized laser lines at excitation wavelengths 836 nm and 1097 nm are used simultaneously thanks to the OPO system. Each scan was acquired with constant laser intensity (20% for 940 nm, 10% for 850 nm, 10% for 800nm, 15% for 836nm, 4% 1097nm) at a 512 × 512 pixel resolution and microscope imaging parameters were maintained over all different regions we imaged. Images, acquired with ZEN software (Zeiss), were saved in .czi format.

### 4.9. Data and statistical analysis

We used ImageJ software to analyze the relative ATP or H_2_O_2_ levels in mitochondria of peripheral axons. The acquired images for each wavelength were aligned using the Template Matching plugin. We defined a Region of Interest (ROI) encompassing all labeled mitochondria of the same axon and the mean fluorescent intensity on the ROI was measured on both images. These light intensities were then corrected for background light intensity determined as an area within the nerve where no fluorescent signal from the viral probe can be observed. Either the ATeam Venus/CFP ratio or the oxidized roGFP/reduced roGFP ratio was then calculated from the 2 values for mean light intensity.

Statistical significance for the effect of time on probe stability was determined using linear regression. The effects of the positive controls and negative controls on probe validation were determined using Student two-tailed T-tests. The effect of nerve stimulation was determined using one-way ANOVA and Dunnett’s post-hoc tests. In addition, the threshold for a responding axon was set at 20% change in probe fluorescence ratio. The differences between MFN2 mutants and wild-type mice were determined using either Student two-tailed T-tests or two-way ANOVA with Sidak post-hoc tests. Differences between internodes and node of Ranvier mitochondria were determined using paired two-tailed T-tests. The effect of demyelination was determined using paired two-tailed T-tests.

### 4.10. Data availability

The data that support the findings of this study are available from the corresponding author upon reasonable request.

## Supporting information

## 5. Acknowledgements

In addition we thank the MRI imaging platform, supported by the French National Research Agency (ANR-10-INBS-04), in particular Hassan Boukhaddaoui, and the Animal Facility of the INM. We also would like to thank Patrice Quintana for his help with the nerve stimulation protocol and Volker Baeker for the image analysis protocol.

This work has benefited from support by an ERC consolidator grant to NT and by the Labex EpiGenMed ANR-10-LABX-12-01.

## 6. Author contributions

GvH conducted the experiments. GvH and NT designed the experiments and wrote the paper with contributions from GC and RC. GC, MD and JB contributed to the experiments and to the design of the experiments. NT supervised the project.

## 10. Expanded view figure legends

**Supplemental Figure 1: LPC injection induces degeneration of the myelin sheath, but not axonal degeneration.**

In a healthy sciatic nerve (Before injection), axons (red) are surrounded by a myelin sheath (green). This myelin sheath is severely damaged following injection of LPC into the sciatic nerve (LPC + 1w), but no sign of axonal degeneration. Injection of PBS (PBS + 1w) does not affect axon or myelin sheath physiology. Scale = 10 μm.

**Supplemental Figure 2: Impact of extramitochondrial ROS on mito-roGFP-Orp1 during demyelination.**

Upon injection of CellROX deep red into the sciatic nerve, fluorescence signal could be detected indicative of oxidative stress. No significant differences were observed between the time points of the demyelination process, nor nerves injected with PBS. In the contralateral nerves, which were not injected with LPC or PBS, a fluorescence signal was detected as well. Before CellROX injection, no fluorescence signal was observed. At the injection site, where tissue is locally damaged due to insertion of the syringe, a much stronger fluorescence signal is observed than at the region where mitochondrial H_2_O_2_ was measured (Fig. 6C).

## References

1. Harris JJ, Jolivet R, Attwell D. Synaptic Energy Use and Supply. Neuron. 2012 Sep 6;75(5):762–77.

2. Attwell D, Laughlin SB. An Energy Budget for Signaling in the Grey Matter of the Brain. J Cereb Blood Flow Metab. 2001 Oct 1;21(10):1133–45.

3. Ames A. CNS energy metabolism as related to function. Brain Res Rev. 2000 Nov 1;34(1):42–68.

4. Salzer JL, Zalc B. Myelination. Curr Biol. 2016 Oct 24;26(20):R971–5.

5. Fischer TD, Dash PK, Liu J, Waxham MN. Morphology of mitochondria in spatially restricted axons revealed by cryo-electron tomography. PLOS Biol. 2018 Sep 17;16(9):e2006169.

6. Misgeld T, Kerschensteiner M, Bareyre FM, Burgess RW, Lichtman JW. Imaging axonal transport of mitochondria in vivo. Nat Methods. 2007 Jun 10;4:559.

7. Zhou B, Yu P, Lin M-Y, Sun T, Chen Y, Sheng Z-H. Facilitation of axon regeneration by enhancing mitochondrial transport and rescuing energy deficits. J Cell Biol. 2016;214(1):103–119.

8. Ohno N, Kidd GJ, Mahad D, Kiryu-Seo S, Avishai A, Komuro H, et al. Myelination and axonal electrical activity modulate the distribution and motility of mitochondria at CNS nodes of Ranvier. J Neurosci Off J Soc Neurosci. 2011 May 18;31(20):7249–58.

9. Perkins GA, Ellisman MH. Mitochondrial Configurations in Peripheral Nerve Suggest Differential ATP Production. J Struct Biol. 2011 Jan;173(1):117–27.

10. Tarasov AI, Griffiths EJ, Rutter GA. Regulation of ATP production by mitochondrial Ca(2+). Cell Calcium. 2012 Jul;52(1):28–35.

11. Edgar JM, McCulloch MC, Thomson CE, Griffiths IR. Distribution of mitochondria along small-diameter myelinated central nervous system axons. J Neurosci Res. 2008;86(10):2250–7.

12. Giorgio M, Trinei M, Migliaccio E, Pelicci PG. Hydrogen peroxide: a metabolic by-product or a common mediator of ageing signals? Nat Rev Mol Cell Biol. 2007 Sep 1;8:722.

13. Murphy MP. How mitochondria produce reactive oxygen species. Biochem J. 2009 Jan 1;417(Pt 1):1–13.

14. Hamanaka RB, Chandel NS. Mitochondrial reactive oxygen species regulate cellular signaling and dictate biological outcomes. Trends Biochem Sci. 2010 Sep;35(9):505–13.

15. Tormos KV, Chandel NS. Seeing the Light: Probing ROS In Vivo Using Redox GFP. Cell Metab. 2011 Dec 7;14(6):720–1.

16. Dixon BJ, Tang J, Zhang JH. The evolution of molecular hydrogen: a noteworthy potential therapy with clinical significance. Med Gas Res. 2013;3:10–10.

17. Tomanek L. Proteomic responses to environmentally induced oxidative stress. J Exp Biol. 2015;218(12):1867–1879.

18. Dauer W, Przedborski S. Parkinson’s disease: mechanisms and models. Neuron. 39th ed. 2003;889–909.

19. Su KG, Banker G, Bourdette D, Forte M. Axonal degeneration in multiple sclerosis: The mitochondrial hypothesis. Curr Neurol Neurosci Rep. 2009 Sep;9(5):411–7.

20. Palau F, Estela A, Pla-Martín D, Sánchez-Piris M. The Role of Mitochondrial Network Dynamics in the Pathogenesis of Charcot-Marie-Tooth Disease. In: Espinós C, Felipo V, Palau F, editors. Inherited Neuromuscular Diseases: Translation from Pathomechanisms to Therapies [Internet]. Dordrecht: Springer Netherlands; 2009. p. 129–37.

21. Smith GM, Gallo G. The role of mitochondria in axon development and regeneration. Dev Neurobiol. 2017 Oct 14;78(3):221–37.

22. Chen H, Chan DC. Emerging functions of mammalian mitochondrial fusion and fission. Hum Mol Genet. 2005 Oct 15;14(suppl_2):R283–9.

23. Misko A, Jiang S, Wegorzewska I, Milbrandt J, Baloh RH. Mitofusin 2 is necessary for transport of axonal mitochondria and interacts with the Miro/Milton complex. J Neurosci Off J Soc Neurosci. 2010 Mar 24;30(12):4232–40.

24. Züchner S, Mersiyanova IV, Muglia M, Bissar-Tadmouri N, Rochelle J, Dadali EL, et al. Mutations in the mitochondrial GTPase mitofusin 2 cause Charcot-Marie-Tooth neuropathy type 2A. Nat Genet. 2004 Apr 4;36:449.

25. Cartoni R, Arnaud E, Médard J-J, Poirot O, Courvoisier DS, Chrast R, et al. Expression of mitofusin 2R94Q in a transgenic mouse leads to Charcot–Marie–Tooth neuropathy type 2A. Brain. 2010 May 1;133(5):1460–9.

26. Pich S, Bach D, Briones P, Liesa M, Camps M, Testar X, et al. The Charcot–Marie–Tooth type 2A gene product, Mfn2, up-regulates fuel oxidation through expression of OXPHOS system. Hum Mol Genet. 2005 Jun 1;14(11):1405–15.

27. Frohman EM, Racke MK, Raine CS. Multiple Sclerosis — The Plaque and Its Pathogenesis. N Engl J Med. 2006 Mar 2;354(9):942–55.

28. Sedel F, Bernard D, Mock DM, Tourbah A. Targeting demyelination and virtual hypoxia with high-dose biotin as a treatment for progressive multiple sclerosis. Oligodendrocytes Health Dis. 2016 Nov 1; 110:644–53.

29. Mahad DJ, Ziabreva I, Campbell G, Lax N, White K, Hanson PS, et al. Mitochondrial changes within axons in multiple sclerosis. Brain J Neurol. 2009 May;132(Pt 5): 1161–74.

30. Campbell G, Mahad DJ. Mitochondrial dysfunction and axon degeneration in progressive multiple sclerosis. FEBS Lett. 2018 Feb 17;592(7):1113–21.

31. Gutscher M, Sobotta MC, Wabnitz GH, Ballikaya S, Meyer AJ, Samstag Y, et al. Proximity-based Protein Thiol Oxidation by H(2)O(2)-scavenging Peroxidases. J Biol Chem. 2009 Nov 13;284(46):31532–40.

32. Imamura H, Huynh Nhat KP, Togawa H, Saito K, Iino R, Kato-Yamada Y, et al. Visualization of ATP levels inside single living cells with fluorescence resonance energy transfer-based genetically encoded indicators. Proc Natl Acad Sci U S A. 2009 Sep 15;106(37): 15651–6.

33. Albrecht SC, Barata AG, Großhans J, Teleman AA, Dick TP. In Vivo Mapping of Hydrogen Peroxide and Oxidized Glutathione Reveals Chemical and Regional Specificity of Redox Homeostasis. Cell Metab. 2011 Dec 7;14(6):819–29.

34. Ren W, Ai H-W. Genetically Encoded Fluorescent Redox Probes. Sensors. 13th ed. 2013;15422–33.

35. Tu H, Boppart SA. Coherent anti-Stokes Raman scattering microscopy: overcoming technical barriers for clinical translation. J Biophotonics. 2014 Jan;7(0):9–22.

36. Mytskaniuk V, Bardin F, Boukhaddaoui H, Rigneault H, Tricaud N. Implementation of a Coherent Anti-Stokes Raman Scattering (CARS) System on a Ti:Sapphire and OPO Laser Based Standard Laser Scanning Microscope. J Vis Exp. 113th ed. 2016;doi: 10.3791/54262.

37. Hajjar H, Boukhaddaoui H, Rizgui A, Sar C, Berthelot J, Perrin-Tricaud C, et al. Label-free non-linear microscopy to measure myelin outcome in a rodent model of Charcot-Marie-Tooth diseases. J Biophotonics. 2018 Aug 9;0(ja):e201800186.

38. Huff TB, Cheng J-X. In vivo coherent anti-Stokes Raman scattering imaging of sciatic nerve tissue. J Microsc. 2007 Feb;225(Pt 2):175–82.

39. Gonzalez S, Fernando R, Berthelot J, Perrin-Tricaud C, Sarzi E, Chrast R, et al. In vivo time-lapse imaging of mitochondria in healthy and diseased peripheral myelin sheath. Mitochondrion. 2015 Jul 1;23:32–41.

40. Ino D, Sagara H, Suzuki J, Kanemaru K, Okubo Y, Iino M. Neuronal Regulation of Schwann Cell Mitochondrial Ca^2+^ Signaling during Myelination. Cell Rep. 2015 Sep 29;12(12):1951–9.

41. Bernard-Marissal N, Van Hameren G, Juneja M, Pellegrino C, Rochat C, El Mansour O, et al. Endoplasmic reticulum and mitochondria dysfunction underlies reversible Charcot–Marie–Tooth type 2A neuropathy.

42. Bazil JN, Beard DA, Vinnakota KC. Catalytic Coupling of Oxidative Phosphorylation, ATP Demand, and Reactive Oxygen Species Generation. Biophys J. 2016 Feb 23;110(4):962–71.

43. Cadenas E, Davies KJA. Mitochondrial free radical generation, oxidative stress, and aging. Free Radic Biol Med. 2000 Aug 1;29(3):222–30.

44. Bélanger E, Henry FP, Vallée R, Randolph MA, Kochevar IE, Winograd JM, et al. In vivo evaluation of demyelination and remyelination in a nerve crush injury model. Biomed Opt Express. 2011 Sep;2(9):2698–2708.

45. Cooper MF, Webster GR. The differentiation of phospholipase A1 and A2 in rat and human nervous tissues. J Neurochem. 1970 Nov;17(11):1543–54.

46. Allt G, Ghabriel MN, Sikri K. Lysophosphatidyl choline-induced demyelination. Acta Neuropathol (Berl). 1988 Sep 1;75(5):456–64.

47. Plemel JR, Michaels NJ, Weishaupt N, Caprariello AV, Keough MB, Rogers JA, et al. Mechanisms of lysophosphatidylcholine-induced demyelination: A primary lipid disrupting myelinopathy. Glia. 2017 Oct 25;66(2):327–47.

48. Kiryu-Seo S, Ohno N, Kidd GJ, Komuro H, Trapp BD. Demyelination increases axonal stationary mitochondrial size and the speed of axonal mitochondrial transport. J Neurosci Off J Soc Neurosci. 2010 May 12;30(19):6658–66.

49. Tricaud N, Park H. Wallerian demyelination: chronicle of a cellular cataclysm. Cell Mol Life Sci. 74th ed. 2017;4049–57.

50. Meiri H, Steinberg R, Medalion B. Detection of Sodium Channel Distribution in Rat Sciatic Nerve Following Lysophosphatidylcholine-Induced Demyelination. J Membrane Biol. 92nd ed. 1986;47–56.

51. Hanson GT, Aggeler R, Oglesbee D, Cannon M, Capaldi RA, Tsien RY, et al. Investigating Mitochondrial Redox Potential with Redox-sensitive Green Fluorescent Protein Indicators. J Biol Chem. 2004 Mar 26;279(13):13044–53.

52. Dooley CT, Dore TM, Hanson GT, Jackson WC, Remington SJ, Tsien RY. Imaging Dynamic Redox Changes in Mammalian Cells with Green Fluorescent Protein Indicators. J Biol Chem. 2004 May 21;279(21):22284–93.

53. Samara C, Poirot O, Domènech-Estévez E, Chrast R. Neuronal activity in the hub of extrasynaptic Schwann cell-axon interactions. Front Cell Neurosci. 2013;7:228.

54. Yi M, Weaver D, Hajnóczky G. Control of mitochondrial motility and distribution by the calcium signal: a homeostatic circuit. J Cell Biol. 2004 Nov 22;167(4):661–72.

55. Fukai T, Ushio-Fukai M. Superoxide Dismutases: Role in Redox Signaling, Vascular Function, and Diseases. Antioxid Redox Signal. 2011 Sep 15;15(6):1583–606.

56. Yamakura F, Kawasaki H. Post-translational modifications of superoxide dismutase. Biochim Biophys Acta. 1804th ed. 2010;318–25.

57. Little C, Olinescu R, Reid KG, O’Brien PJ. Properties and Regulation of Glutathione Peroxidase. J Biol Chem. 1970 Jul 25;245(14):3632–6.

58. Nie Q, Wang C, Song G, Ma H, Kong D, Zhang X, et al. Mitofusin 2 deficiency leads to oxidative stress that contributes to insulin resistance in rat skeletal muscle cells. Mol Biol Rep. 2014 Oct 1;41(10):6975–83.

59. Baloh RH, Schmidt RE, Pestronk A, Milbrandt J. Altered Axonal Mitochondrial Transport in the Pathogenesis of Charcot-Marie-Tooth Disease from Mitofusin 2 Mutations. J Neurosci. 2007;27(2):422–430.

60. Zilberter Y, Zilberter T, Bregestovski P. Neuronal activity in vitro and the in vivo reality: the role of energy homeostasis. Trends Pharmacol Sci. 2010 Sep 1;31(9):394–401.

61. Ainbinder A, Boncompagni S, Protasi F, Dirksen RT. Role of Mitofusin-2 in Mitochondrial Localization and Calcium Uptake in Skeletal Muscle. Cell Calcium. 2015 Jan;57(1):14–24.

62. Chen Y, Csordás G, Jowdy C, Schneider TG, Csordás N, Wang W, et al. Mitofusin 2-containing Mitochondrial-Reticular Microdomains Direct Rapid Cardiomyocyte Bioenergetic Responses via Inter-Organelle Ca(2+) Crosstalk. Circ Res. 2012 Sep 14;111(7):863–75.

63. Yu T, Robotham JL, Yoon Y. Increased production of reactive oxygen species in hyperglycemic conditions requires dynamic change of mitochondrial morphology. Proc Natl Acad Sci U S A. 2006 Feb 21;103(8):2653–8.

64. de Brito OM, Scorrano L. Mitofusin 2: A Mitochondria-Shaping Protein with Signaling Roles Beyond Fusion. Antioxid Redox Signal. 2007 Dec 20;10(3):621–34.

65. Giorgio M, Migliaccio E, Orsini F, Paolucci D, Moroni M, Contursi C, et al. Electron Transfer between Cytochrome c and p66Shc Generates Reactive Oxygen Species that Trigger Mitochondrial Apoptosis. Cell. 2005 Jul 29;122(2):221–33.

66. Chen H, Chan DC. Mitochondrial dynamics–fusion, fission, movement, and mitophagy–in neurodegenerative diseases. Hum Mol Genet. 2009 Oct 15;18(R2):R169–76.

67. Bros H, Millward JM, Paul F, Niesner R, Infante-Duarte C. Oxidative damage to mitochondria at the nodes of Ranvier precedes axon degeneration in ex vivo transected axons. Exp Neurol. 2014 Nov 1;261:127–35.

68. Witte ME, Bø L, Rodenburg RJ, Belien JA, Musters R, Hazes T, et al. Enhanced number and activity of mitochondria in multiple sclerosis lesions. J Pathol. 2009 May 26;219(2):193–204.

69. Young EA, Fowler CD, Kidd GJ, Chang A, Rudick R, Fisher E, et al. Imaging correlates of decreased axonal Na+/K+ ATPase in chronic multiple sclerosis lesions. Ann Neurol. 2008;63(4):428–35.

70. Gilgun-Sherki Y, Melamed E, Offen D. The role of oxidative stress in thepathogenesis of multiple sclerosis: The need for effectiveantioxidant therapy. J Neurol. 2004 Mar 1;251(3):261–8.

71. Rousset S, Alves-Guerra M-C, Mozo J, Miroux B, Cassard-Doulcier A-M, Bouillaud F, et al. The Biology of Mitochondrial Uncoupling Proteins. Diabetes. 2004;53(suppl 1):S130–S135.

72. Dalgaard LT, Pedersen O. Uncoupling proteins: functional characteristics and role in the pathogenesis of obesity and Type II diabetes. Diabetologia. 2001 Aug 1;44(8):946–65.

73. Rexroth S, Poetsch A, Rögner M, Hamann A, Werner A, Osiewacz HD, et al. Reactive oxygen species target specific tryptophan site in the mitochondrial ATP synthase. Biochim Biophys Acta BBA - Bioenerg. 2012 Feb 1;1817(2):381–7.

74. Yin X, Kidd GJ, Ohno N, Perkins GA, Ellisman MH, Bastian C, et al. Proteolipid protein–deficient myelin promotes axonal mitochondrial dysfunction via altered metabolic coupling. J Cell Biol. 2016 Nov 21;215(4):531–42.

75. Yellen G. Fueling thought: Management of glycolysis and oxidative phosphorylation in neuronal metabolism. J Cell Biol. 2018;217(7):2235–2246.

76. Funfschilling U, Supplie LM, Mahad D, Boretius S, Saab AS, Edgar J, et al. Glycolytic oligodendrocytes maintain myelin and long-term axonal integrity. Nature. 2012 Apr 29;485(7399):517–21.

77. Stefanini M, Martino CD, Zamboni L. Fixation of Ejaculated Spermatozoa for Electron Microscopy. Nature. 1967 Oct 14;216:173.

78. Hung H, Kohnken R, Svaren J. The NURD chromatin remodeling complex is required for peripheral nerve myelination. J Neurosci. 2012 Feb 1;32(5):1517–27.

