## Supplementary figures and images for "CMT disease 2A and demyelination decouple ATP and ROS production by axonal mitochondria"

### Supplementary file 1

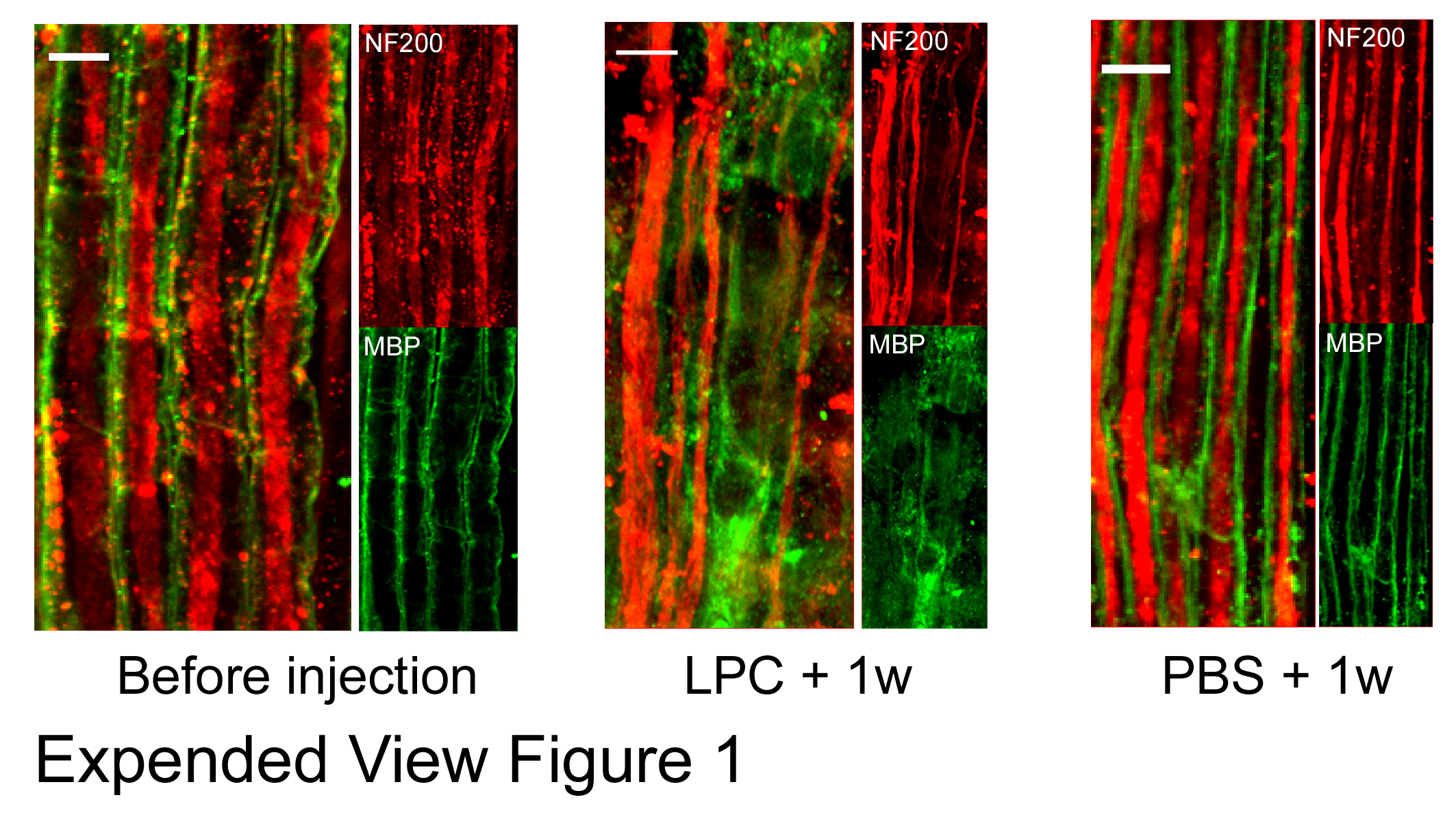

### Supplementary file 2

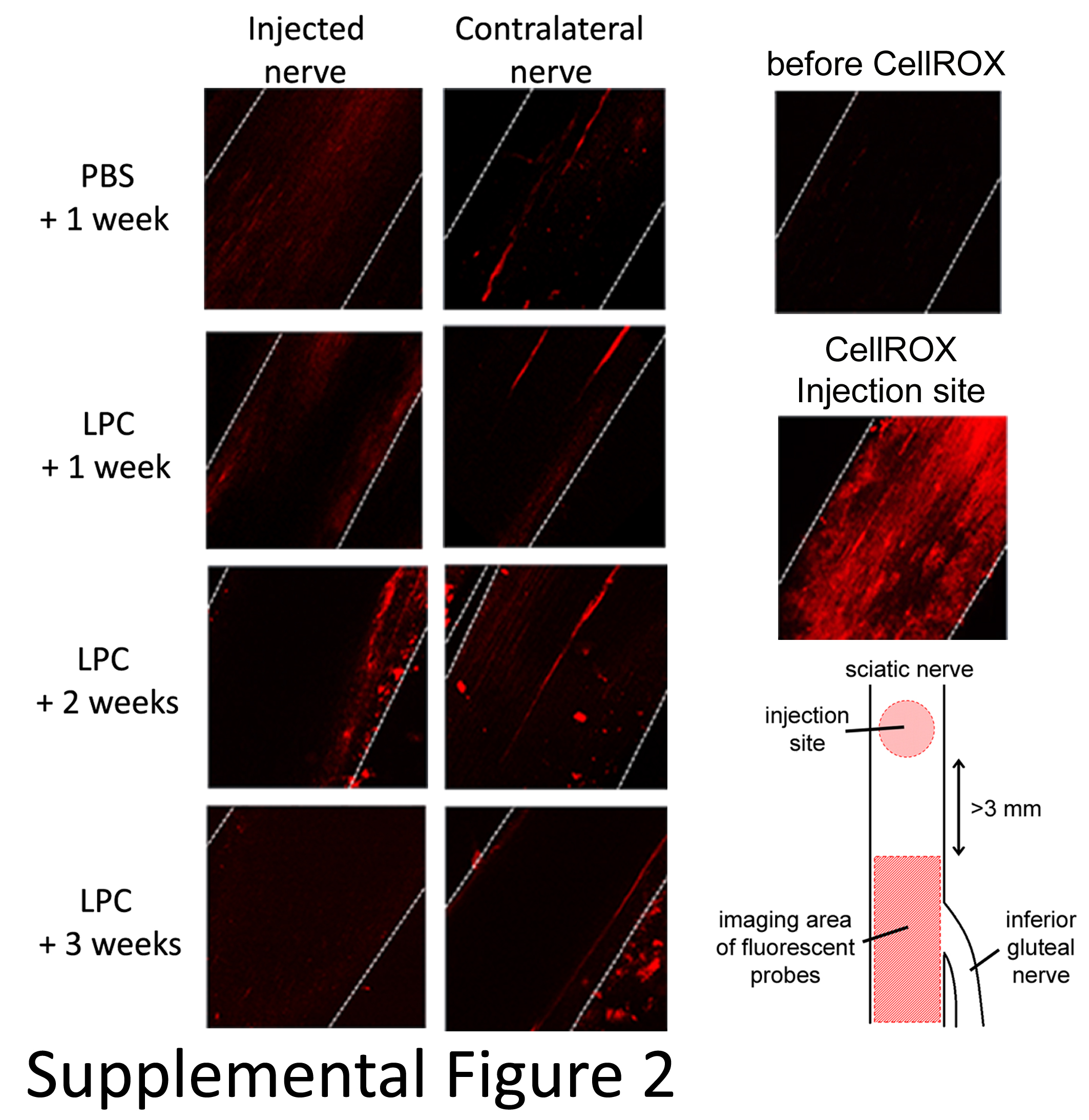
